# Order under uncertainty: robust differential expression analysis using probabilistic models for pseudotime inference

**DOI:** 10.1101/047365

**Authors:** Kieran Campbell, Christopher Yau

## Abstract

Single cell gene expression profiling can be used to quantify transcriptional dynamics in temporal processes, such as cell differentiation, using computational methods to label each cell with a ‘pseudotime’ where true time series experimentation is too difficult to perform. However, owing to the high variability in gene expression between individual cells, there is an inherent uncertainty in the precise temporal ordering of the cells. Preexisting methods for pseudotime ordering have predominantly given point estimates precluding a rigorous analysis of the implications of uncertainty. We use probabilistic modelling techniques to quantify pseudotime uncertainty and propagate this into downstream differential expression analysis. We demonstrate that reliance on a point estimate of pseudotime can lead to inflated false discovery rates compared and that probabilistic approaches provide greater robustness and measures of the temporal resolution that can be obtained from pseudotime inference.

## Background

The emergence of high-throughput single cell genomics as a tool for the precision study of biological systems Kalisky and Quake (2011); Shapiro et al. (2013); Macaulay and Voet (2014); Wills and Mead (2015) has given rise to a variety of novel computational and statistical modelling challenges Stegle et al. (2015); Trapnell (2015). One particular area of interest has been the study of transcriptional dynamics in temporal processes, such as cell differentiation or proliferation Treutlein et al. (2014); Tsang et al. (2015), in order to understand the coordinated changes in transcription programming that underlie these processes. In the study of such systems, practical experimental designs that can allow the collection of real time series data maybe difficult or impossible to achieve. Instead, investigators have adopted computational methods to identify temporal signatures and trends from unordered genomic profiles of single cells, a process known as *pseudotemporal ordering* Qiu et al. (2011); Bendall et al. (2014); Marco et al. (2014); Trapnell et al. (2014); Moignard et al. (2015); Reid and Wernisch (2015). Computational approaches for this problem were first tackled in the context of gene expression microarray analysis of bulk cell populations Magwene et al. (2003); Gupta and Bar-Joseph (2008); Qiu et al. (2011) but the recent availability of single cell technology overcomes the limitations of measuring population averaged signals in bulk analyses.

Pseudotemporal ordering of whole-transcriptome profiles of single cells with unsupervised computational methods has an advantage over cytometry-based assays in that it does not rely on *a priori* knowledge of marker genes. The principle underlying these methods is that each single cell RNA sequencing experiment constitutes a time series in which each cell represents a distinct time point along a continuum representing the underlying degree of temporal progress (Figure 1A). During the single cell capture process, the *true* temporal label that identifies the stage of the cell is lost (Figure 1B) and these parameters become latent, unobserved quantities that must be statistically inferred from the collection of single cell expression profiles (Figure 1C). Importantly, absolute physical time information will in general be irretrievably lost and it is only possible to assign a “pseudotime” for each cell that provides a relative quantitative measure of progression. Consequently, whilst the correspondence between physical and pseudotime ordering maybe conserved, the pseudotimes themselves are not necessarily calibrated to actual physical times. Pseudotime ordering can potentially be used to recapitulate temporal resolution in an experiment that does not explicitly capture labelled time series data. The pseudotimes could then be used to identify genes that are differentially expressed across pseudotime (Figure 1D) providing insight into the evolution of transcription programming.

**Figure 1:**
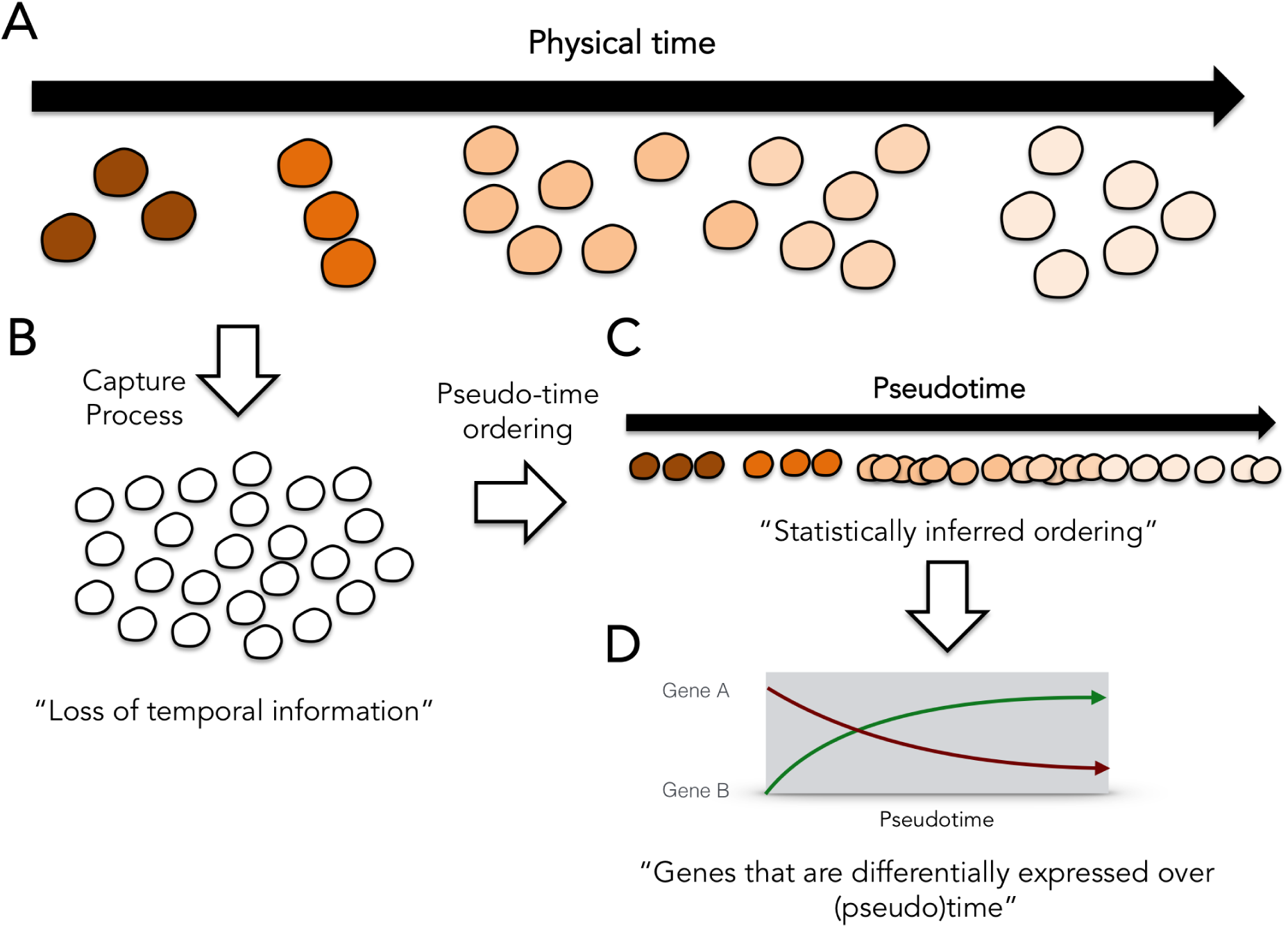
The single cell pseudotime ordering problem. (A) Single cells at different stages of a temporal process. (B) The temporal labelling information is lost during single cell capture. (C) Statistical pseudotime ordering algorithms attempt to reconstruct the relative temporal ordering of the cells but cannot fully reproduce physical time. (D) The pseudotime estimates can be used to identify genes that are differentially expressed over (pseudo)time.

Practically, current methods for pseudotime inference proceed via a multi-step process. First, gene selection and dimensionality reduction techniques are applied to compress the infor-mation held in the high-dimensional gene expression profiles to a small number of dimensions (typically two or three for simplicity of visualisation). The identification of an appropriate dimensionality reduction technique is a *subjective* choice and a number of methods have been adopted such as Principal and Independent Components Analysis (P/ICA) and highly nonlinear techniques such as diffusion maps Haghverdi et al. (2015) or stochastic neighbourhood embedding (SNE) Hinton and Roweis (2002); Van der Maaten and Hinton (2008); Amir et al. (2013). This choice is guided by whether the dimensionality reduction procedure is able to identify a suitable low-dimensional embedding of the data that contains a relatively smooth trajectory that might plausibly correspond to the temporal process under investigation.

Next, the pseudotime trajectory of the cells in this low-dimensional embedding is characterised. In Monocle Trapnell et al. (2014) this is achieved by the construction of a minimum spanning tree (MST) joining all cells. The diameter of the MST provides the main trajectory along which pseudotime is measured. Related graph-based techniques (Wanderlust) have also been used to characterise temporal processes from single cell mass cytometry data Bendall et al. (2014). In SCUBA Marco et al. (2014) the trajectory itself is directly modelled using principal curves Hastie and Stuetzle (2012) and pseudotime is assigned to each cell by projecting its location in the low-dimensional embedding on to the principal curve. The estimated pseudo-times can then be used to order the cells and to assess differential expression of genes across pseudotime.

A limitation of these approaches is that they provide only a single *point estimate* of pseudotimes concealing the full impact of gene expression variability and technical noise. As a consequence, the statistical uncertainty in the pseudotimes is not propagated to downstream analyses precluding a thorough treatment of robustness and stability. To date, the impact of this pseudotime uncertainty has not been explored and its implications are unknown as the methods applied do not possess a probabilistic interpretation. However, we can examine the stability of the pseudotime estimates by taking multiple random subsets of a dataset and re-estimating the pseudotimes for each subset. For example, we have found that the pseudotime assigned to the same cell can vary considerably across random subsets in Monocle (details given in Supplementary Materials and Supplementary Figure S1).

In order to address pseudotime uncertainty in a formal and coherent framework, probabilistic approaches using Gaussian Process Latent Variable Models (GPLVM) have been used recently as non-parametric models of pseudotime trajectories Reid and Wernisch (2015); Campbell and Yau (2015). These provide an explicit model of pseudotimes as latent embedded one-dimensional variables and can be fitted within a Bayesian statistical framework allowing posterior uncertainty in the pseudotimes to be derived using Markov Chain Monte Carlo (MCMC) simulations. In this article we adopt this framework based to assess the impact of pseudotime uncertainty on downstream differential analyses. We will show that pseudotime uncertainty can be non-negligible and when propagated to downstream analysis may considerably inflate false discovery rates. We demonstrate that there exists a limit to the degree of recoverable temporal resolution, due to intrinsic variability in the data, with which we can make statements such as “this cell precedes another”. Finally, we propose a simple means of accounting for the different possible choices of reduced dimension data embeddings. We demonstrate that, given sensible choices of low-dimensional representations, these can be combined to produce more robust pseudotime estimates. Overall, we outline a modelling and analytical strategy to produce more stable pseudotime based differential expression analysis.

## Results

### Probabilistic pseudotime inference using Gaussian Process Latent Variable Models

We first provide a brief overview of the Gaussian Process Latent Variable Model Titsias and Lawrence (2010). The GPLVM uses a Gaussian Process to define a stochastic mapping between a low-dimensional latent space to a (typically) higher dimensional observation space. A Gaussian Process is characterised by a mean function describing the expected mapping between the latent and observation spaces and a covariance function that describes the covariance between the mapping function evaluated at any two arbitrary latent positions. The covariance function therefore acts to control the second-order statistics of the Gaussian Process and suitable choices can be designed to encourage properties such as smoothness, periodicity or other second-order features.

For this application, the latent space is one-dimensional, describing pseudotime progression whilst the observations are the reduced dimensionality representations of the single cell expres-sion data. We will use Bayesian inference to characterise the joint posterior distribution p(t|X) of the pseudotimes **t** = {*t*_1_,…, *t*_*n*_} given the expression data **X** = {x_1_,…, x_n_} for *n* single cells. As the integrals involved are mathematically intractable, we will use Markov Chain Monte Carlo simulations to obtain a numerical approximation to the posterior by drawing samples from the posterior distribution. Each sample corresponds to one possible trajectory *and* ordering for the cells with the set of samples providing an approximate distribution of pseudotimes. The pseudotime values are between measured 0 and 1 where a value of 0 corresponds to one end state of the temporal process and a value 1 to the other. In this work we focus only on non-bifurcating processes. Figure 2 gives a diagrammatic representation of our proposed workflow and a more detailed model descriptions is given in Methods.

**Figure 2:**
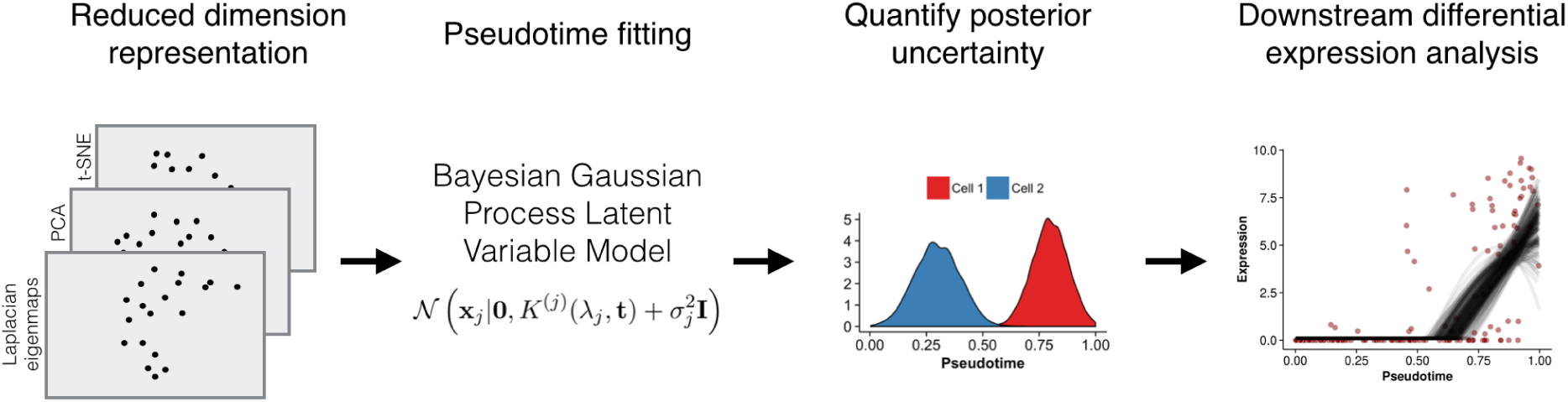
Workflow for fitting Bayesian Gaussian Process Latent Variable Model pseudotime models. Reduced-dimension representations of the gene expression data (from Laplacian eigenmaps, PCA and/or t-SNE) are created. The pseudotime can be fitted using one or more low dimensional representations of the data. Posterior samples of pseudotimes are drawn from a Bayesian GPLVM and these are used to obtain alternative pseudotime estimates. Downstream differential analyses can be performed on the posterior samples to characterise the robustness with respect to variation in pseudotime ordering.

### Sources of uncertainty in pseudotime inference

We applied our probabilistic pseudotime inference to three published single-cell RNA-seq datasets of differentiating cells: myoblasts in Trapnell et al. (2014) Trapnell et al. (2014), hippocampal quiescent neural stem cells in Shin et al. (2015) Shin et al. (2015) and sensory epithelia from the inner ear in Burns et al. (2015) Burns et al. (2015). For the Trapnell and Shin datasets we used Laplacian Eigenmaps Belkin and Niyogi (2003) for dimensionality reduction prior to pseudotime inference, while for the Burns dataset we used the PCA representation of the cells from the original publication (for a detailed description of our analysis, see Supplementary Methods: Data Analysis). These particular choices of reduced dimensionality representations gave visually plausible trajectory paths in two dimensions.

An implicit assumption in pseudotime ordering is that proximity in pseudotime should reflect proximity in the observation or data space. That is, two cells with similar pseudotime assignments should have similar gene expression profiles but, in practice, cell-to-cell variability and technical noise means that the location of the cells in the observation space will be variable even if they truly do have the same pseudotime. We plotted posterior mean pseudotime trajectories for the three datasets learned using the GPLVM in Figure 3A-C and the posterior predictive data distribution *p*(**X*|X**). The posterior predictive data distribution gives an indication of where *future* data points might occur given the existing data. Notice that for all three data sets, this distribution can be quite diffuse. This is due to a combination of actual cell-to-cell expression variability, manifesting as a spread of data points around the mean trajectory, but also model misspecification (the difference between what our “assumed” model and the “true” but unknown data generating mechanism).

**Figure 3:**
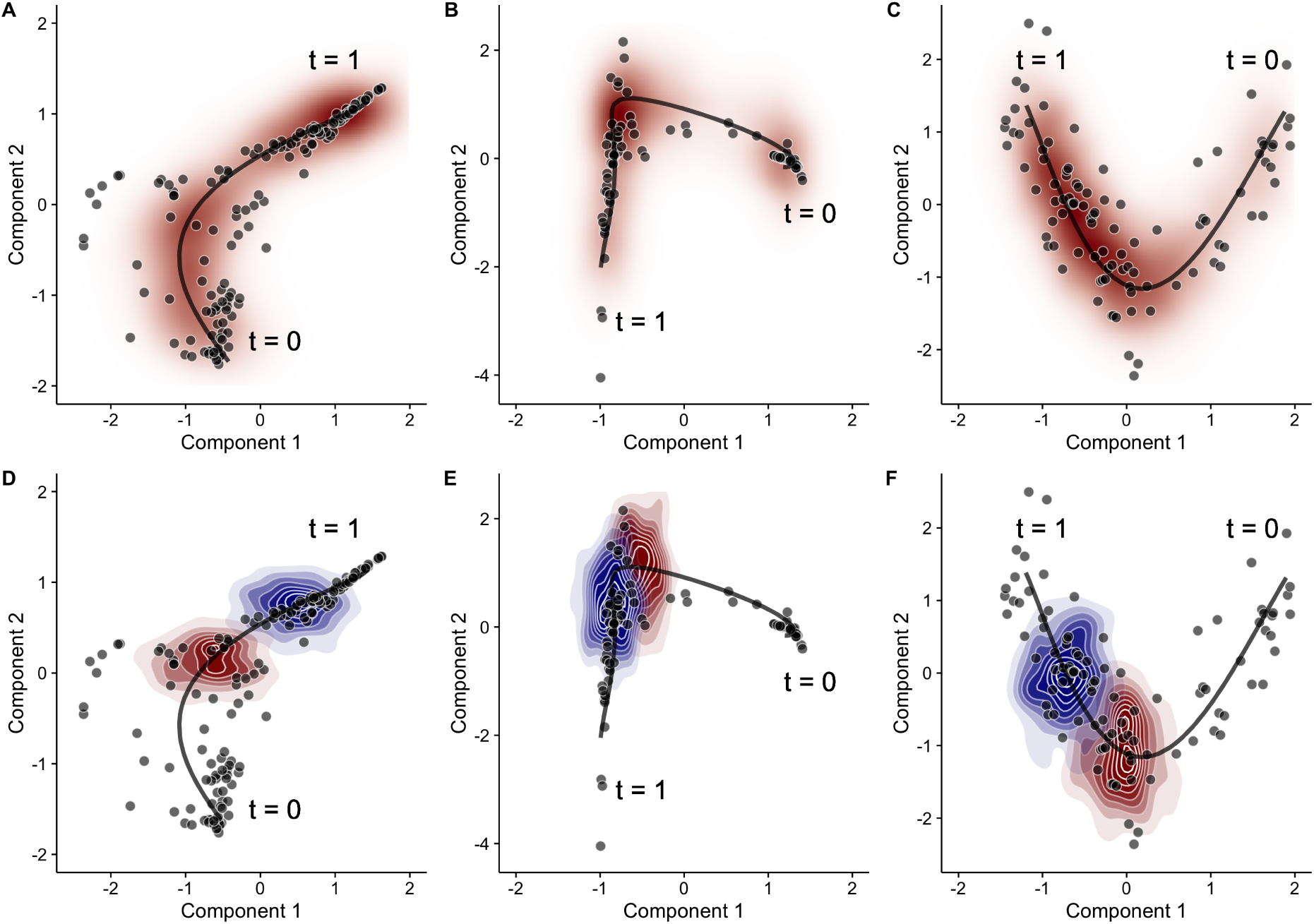
Posterior pseudotime trajectories for three single-cell RNA-seq datasets. Posterior pseudotime trajectories shown in a two-dimensional reduced representation space for (left) a Laplacian eigenmaps representation of Trapnell et al. (2014) Trapnell et al. (2014), (centre) Laplacian eigenmaps representation of Burns et al. (2015) Burns et al. (2015) and (right) PCA representation of Shin et al. (2015) Shin et al. (2015). Each point represents a cell and the black line represents the mean pseudotime trajectory. Plots (**A-C**) shows the overall posterior predictive data density (red) whilst (**D-F**) shows the conditional posterior predictive data density for *t* = 0.5 (red) and *t* = 0.7 (blue).

It is interesting to discuss the latter point as it is an issue that is often not adequately addressed or fully acknowledged in the literature. The GPLVM applied assumes a homoscedastic noise distribution which is uniform along the pseudotime trajectory. However, it is clear that the variability of the data points can change along the trajectory and a heteroscedastic (non-uniform) noise model may be more appropriate in certain scenarios. Unfortunately, whilst models of heteroscedastic noise processes can be applied Le et al. (2005), these typically severely complicate the statistical inference and require a model of how the variability changes over pseudotime which is likely to be unknown. The important point here is that the posterior probability calculations are always calibrated with respect to a given model. The better the model represents the true data generating mechanism, the better calibrated the probabilities. Model misspecification can also contribute to posterior uncertainty in inferred parameters.

Returning to the intrinsic cell-to-cell variability, we next considered the conditional posterior predictive data distributions *p*(**X**^*^|*t*^*^, **X**) which are shown in Figure 3D-F. These distributions show the possible distribution of future data points given the existing data and a theoretical pseudotime *t*^*^ and, in this example, we condition on pseudotimes *t*^*^ = 0.5 and *t*^*^ = 0.7. Although the two pseudotimes differ by a magnitude of 0.2, the conditional predictive distributions are very close or overlapping. This means that cells with pseudotimes of 0.5 or 0.7 could have given rise to data point occupying these overlapping regions. This variability is what ultimately limits the temporal resolution that can be obtained.

It is important to note that the posterior mean trajectories correspond to certain a priori or subjective smoothness assumptions (specified as hyperparameters in the model specification) which dedicate the curvature properties of the trajectory. Figure 4 shows three alternative posterior mean pseudotime trajectories for the Trapnell data based on different hyperparameters settings for the GPLVM. In a truly unsupervised scenario all three paths could be plausible as we would have little information to inform us about the true shape of the trajectory. This would become an additional source of uncertainty in the pseudotime estimates. However, we favoured hyperparameter settings that gave rise to well-defined (unimodal) posterior distributions that resulted in multiple independent Markov Chain Monte Carlo runs converging to the same mean trajectory rather than settings that give rise to a “lumpy” posterior distribution with many local modes corresponding to different interpretations of the data (see Supplementary Figure S3). Later on, when we consider inference using multiple representations, the ability to specify a wider choice of trajectories is useful as we will demonstrate how the correspondence between pseudotime trajectories in different reduced dimension representations is not always obvious from a visual analysis.

**Figure 4:**
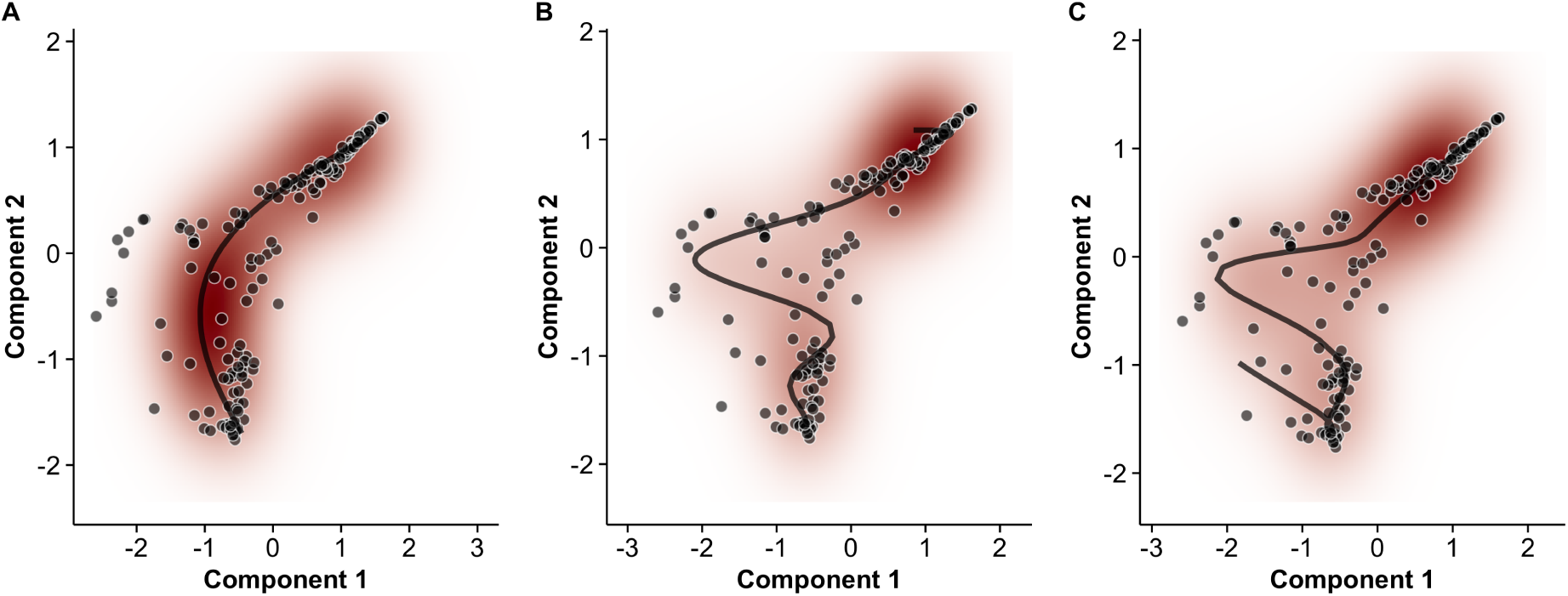
Effect of prior expectations on pseudotime trajectories. The prior probability distribution (defined in terms of hyperparameters (*γ*_α_, γ_β_) in our model) on the expected smoothness of pseudotime trajectories can fundamentally change the inferred progression path. Examples shown using the data of Trapnell et al. (2014) Trapnell et al. (2014). Red - shows the density of the posterior predictive data distribution. Black - shows the mean pseudotime trajectory. Shrinkage hyperparameters (γ_α_, γ_β_) of (30, 5), (5,1) and (3,1) were used for A, B and C respectively.

We next examined the posterior distributions in pseudotime assignment for four cells from the Trapnell dataset in Figure 5A. Uncertainty in the estimate of pseudotime is assessed using the highest probability density (HPD) credible interval (CI), the Bayesian equivalent of the confidence interval. The 95% pseudotime CI typically covers around one quarter of the trajectory, suggesting that pseudotemporal orderings of single-cells can potentially only resolve a cell’s place within a trajectory to a coarse estimate (e.g. ‘beginning’, ‘middle’ or ‘end’) and do not necessarily dramatically increase the temporal resolution of the data. One immediate consequence of this is that it is unlikely that we can make definite statements such as whether one cell comes exactly before or after another. This is illustrated in Figures 5B-D which displays the estimated pseudotime uncertainty for all three datasets. In all the datasets, the general progression is apparent, but the precise ordering of the cells has a non-trivial degree of ambiguity.

**Figure 5:**
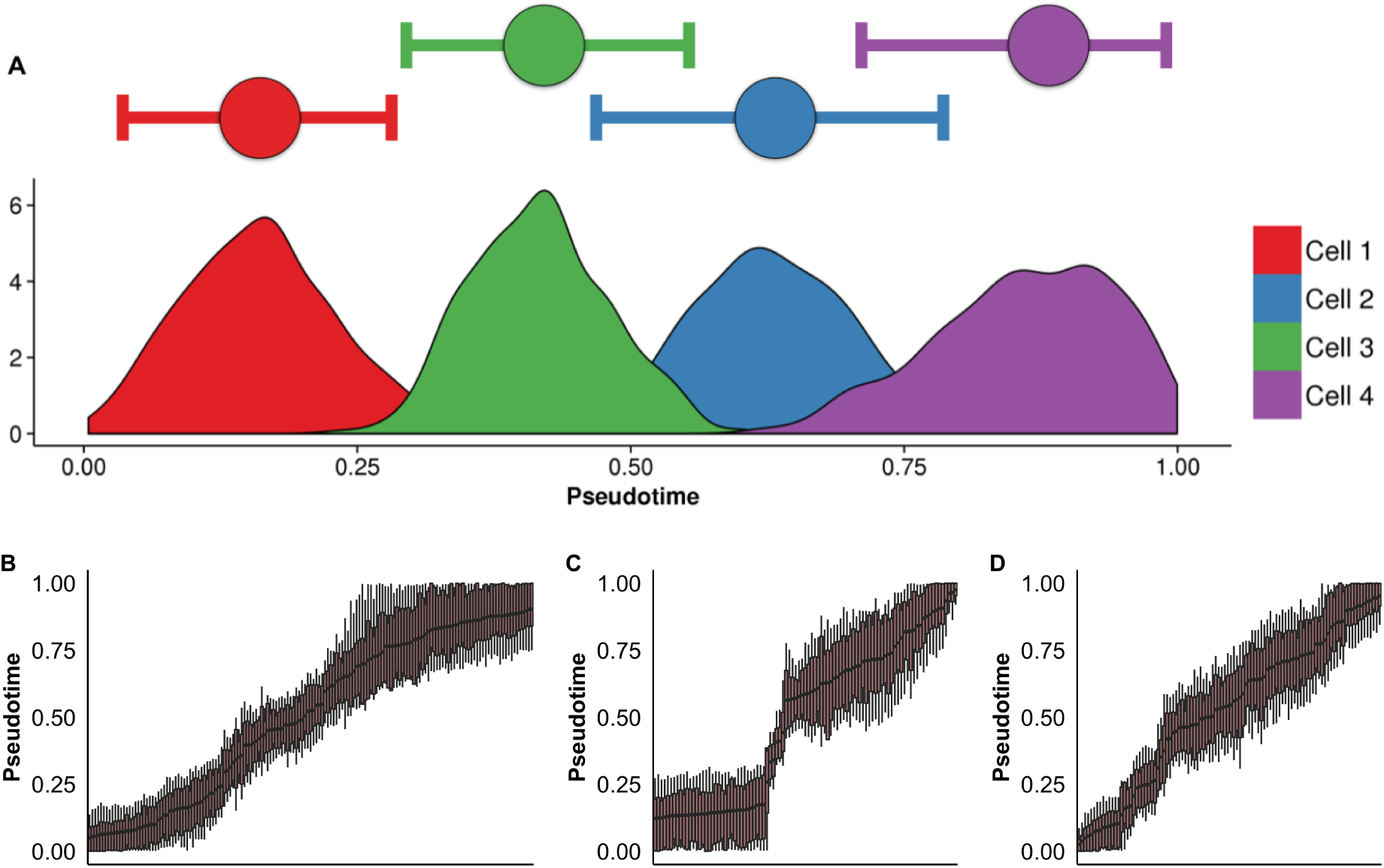
Posterior uncertainty in pseudotime trajectories. (A) Posterior uncertainty in pseudotimes for four randomly selected cells from the Trapnell et al. (2014) dataset. Horizontal bars represent the 95% highest probability density (HPD) credible interval (CI), which typically covers around a quarter of the pseudotime trajectory. (B-D) Boxplots showing the posterior uncertainty for each cell from the Trapnell et al. (2014) datasets. The edges of the boxes and tails correspond to the 75% and 95% HPD-CIs respectively.

### Failure to account for pseudotime uncertainty leads to increased false discovery rates

The previous section addressed the sources of statistical uncertainty in the pseudotimes. We next explored the impact of pseudotime uncertainty on downstream analysis. Specifically, we focused on the identification of genes that are differentially expressed across pseudotime. Typically, these analyses involve regression models that assume the input variables (the pseudotimes) are both fixed and certain but, with our probabilistic model, we can use the posterior samples from our Bayesian model to refit the regression model to each pseudotime estimate. In doing so we can examine which genes are called as significant in each of the posterior samples and assess the stability of the differential expression analysis to pseudotime uncertainty by recording how frequently genes are designated as significant across the posterior samples. This allowed us to re-estimate the false discovery rate (FDR) fully accounting for the variability in pseudotime. As there are a multitude of sources of uncertainty on top of this (such as biological and technical variability) this allows us to put a lower bound on the FDR of such analyses in general.

Precisely, we fitted the tobit regression model from Trapnell et al. (2014) for each gene for each sample from the posterior pseudotime distribution, giving us a per-gene set of false-discovery-rate-corrected *Q*-values. We then compared the proportion of times a gene is called as differentially expressed (5% FDR) across all pseudotime samples to the *Q*-value using a point pseudotime estimate based on the maximum a *posteriori* (or MAP) estimate. We reasoned that if a gene is truly differentially expressed then such expression will be robust to the underlying uncertainty in the ordering. Note for comparison, our MAP estimates with the GPLVM correlate strongly with Monocle derived pseudotime point estimates (see Supplementary Figure S2).

Figure 6A shows two analyses for two illustrative genes (ITGAE and ID2) in the Trapnell data set. Using the MAP pseudotime ordering, differential expression analysis of ITGAE over pseudotime attained a q-value of 0.02. However, the gene was only called significant in only 9% of posterior pseudotime samples with a median q-value of 0.32. In contrast, ID1 - known to be involved in muscle differentiation - had a *q*-value of 6.6 × 10^−11^ using the MAP pseudotime ordering, but was also called significant in all the posterior pseudotime ordering samples having a median q-value of 4.4 × 10^−11^. This indicates that the significance of the temporal expression variability of ID1 is robust with respect to posterior sampling of the pseudotime ordering whilst the significance ITGAE is much more dependent on the ordering chosen.

**Figure 6:**
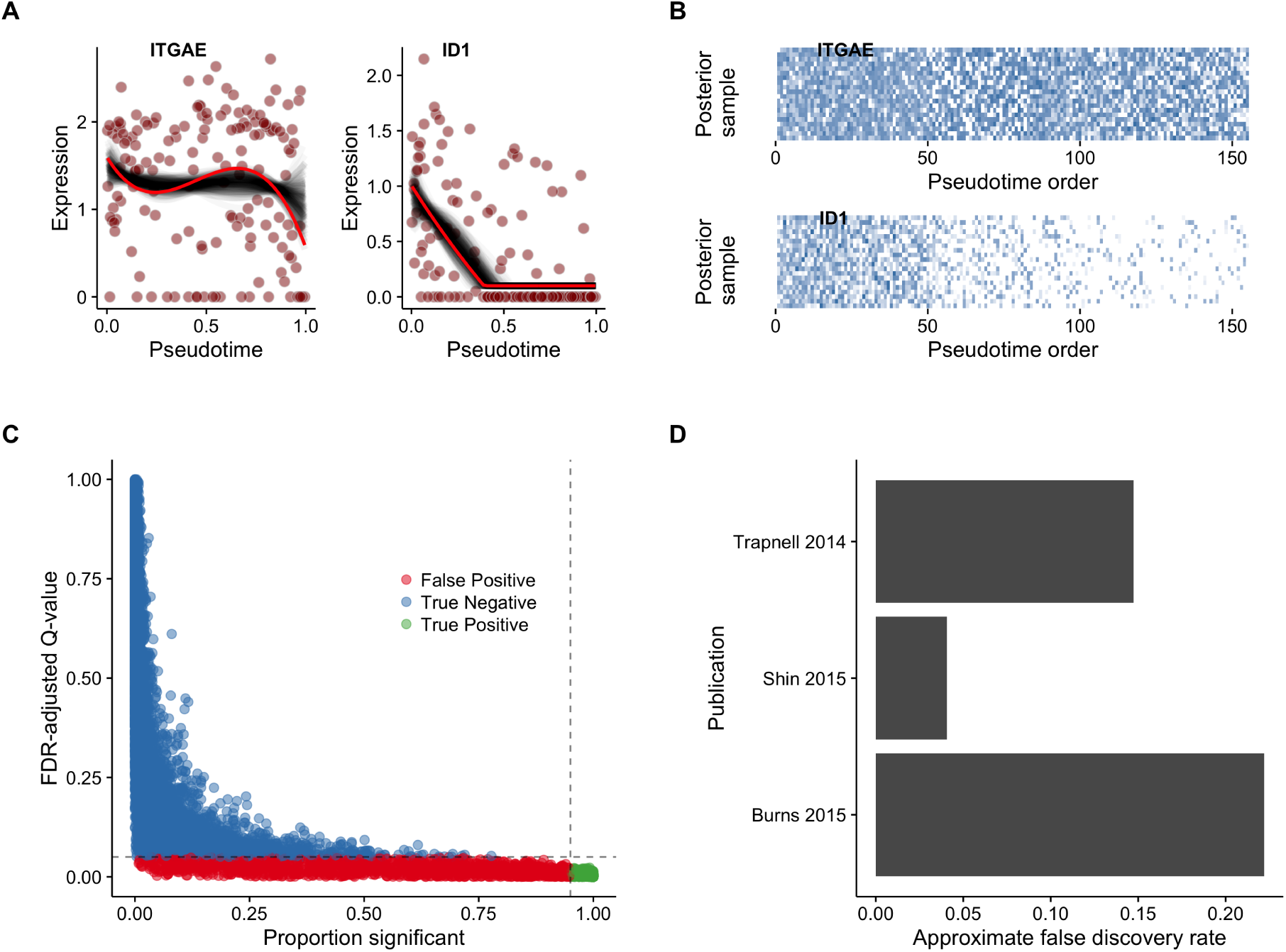
Approximate FDR for differential expression across pseudotime. (**A**) Gene expression plots across pseudotime, with black traces corresponding to models fitted to pseudotime samples while the red trace corresponds to the point (MAP) estimate for two exemplar genes and (**B**) corresponding posterior pseudotime orderings. (**C**) Scatter plot of point estimate q-values against proportion significant for all genes (Trapnell dataset). (**D**) Approximate false discovery rates (AFDR) for three datasets (Trapnell et al. 2014, Shin et al. 2015 and Burns et al. 2015).

As a conservative rule of thumb, we designated a putative temporal association as a false positive if the gene has a *Q*-value less than 5% at the MAP estimate of pseudotime but is significant in less than 95% of the posterior pseudotime samples. Looking across all genes in the Trapnell data, Figure 6B shows that a significant number of genes that were found to have a *Q*-value < 0.05 and deemed significant based on the MAP pseudotime ordering, did not replicate consistently and were not robust to alternate orderings. In fact, across the three datasets we analysed, we found that the false discovery rate, when adjusted for pseudotime uncertainty, ranged from 4% to 20% (Figure 6C). This indicates the FDR can be up to much larger than the expected 5% and crucially is variable across datasets meaning there is no simple rule of thumb that can be applied a priori to account for pseudotime uncertainty. Such values remain low enough that analyses examining the coexpression of gene sets across pseudotime (such as in Trapnell et al. (2014)) will still be largely valid. However, if a set of robustly differentially expressed genes is required or the FDR needs to be characterised then a full probabilistic treatment of pseudotime is needed.

### A sigmoidal model of switch-like behaviour across pseudotime

In the previous section, we examined differential expression across pseudotime by fitting generalized additive models to the gene expression profiles Trapnell et al. (2014). Their approach used a Tobit regression model with a cubic smoothing spline link function. Hypothesis testing using the likelihood ratio test is conducted against a null model of no pseudotime dependence. This model provides a highly flexible but non-specific model of pseudotime dependence that was not suited to the next question we wished to address.

Specifically, we were interested in whether we could identify if two genes switched behaviours at the same (or similar) times during the temporal process and therefore an estimate of the time resolution that can be gained from a pseudotime ordering approach. This requires estimation of a parameter that can be directly linked to a switch on(/off) time that is not present in the Tobit regression model. As a result, we propose a “sigmoidal” model of differential expression across pseudotime that better captures switch-like gene (in)activation and has easy to interpret parameters corresponding to activation time and strength. By combining such a parametric model with the Bayesian inference of pseudotime we can then infer the resolution to which we can say whether one gene switches on or off before another. Details of the sigmoidal gene activation model are given in Methods and in Supplementary Methods.

We applied our sigmoidal model to learn patterns of switch-like behaviour of genes in the Trapnell dataset. For each gene we estimated the *activation time* (*t*_0_) as well as the *activation strength* (*k*). We fitted these sigmoidal switching models to all posterior pseudotime samples to approximate the posterior distribution for the time and strength parameters. We uncovered a small set of genes whose median activation strength is distinctly larger than the rest and had low variability across posterior pseudotime samples implying a population of genes that exhibit highly switch-like behaviour (Figure 7A). Some genes showed high activation strength for certain pseudotime orderings but low overall median levels across all the posterior samples. We concluded that genes with large credible intervals on the estimates of activation strength do not show robust switch-like behaviour and demonstrate the necessity of using probabilistic methods to infer gene behaviour as opposed to point estimates that might give highly unstable results.

**Figure 7:**
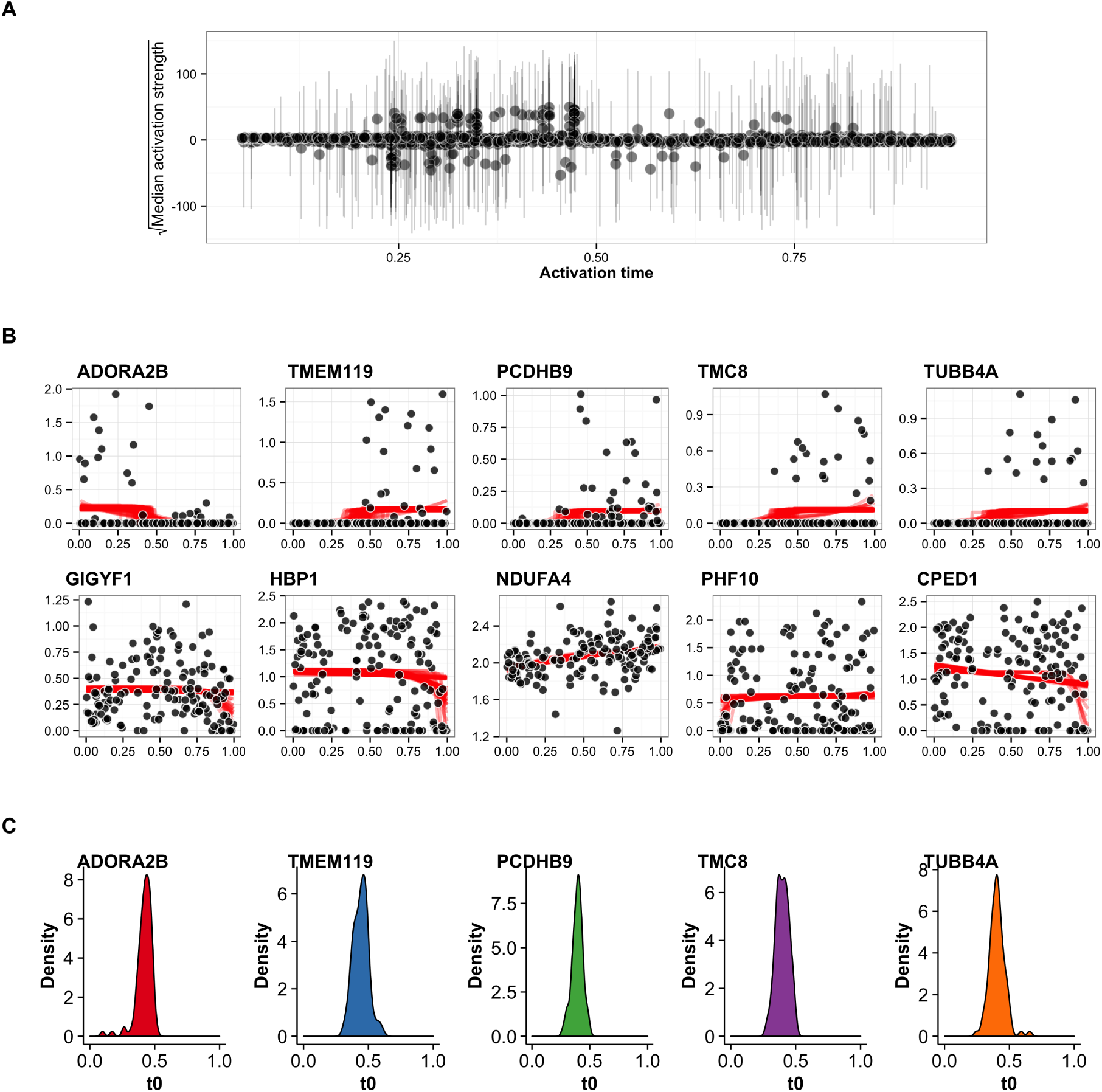
Robust inference of switch-like behaviour in genes across pseudotime. (**A**) The square-root of the median of the activation strength parameter *k* across all pseudotime samples as a function of activation time *t*_0_. The error bars show the 95% credible interval, demonstrating that point estimates can severely skew the apparent behaviour of genes and a requirement for a robust Bayesian treatment of gene expression. A distinct population of genes whose median activation strength sits separate from the majority close to the x-axis implies a subset of genes show true switch-like behaviour. (**B**) Representative examples of genes whose median activation strength is large (top row) compared to small (bottom row). Each black point represents the gene expression of the cell with red lines corresponding to posterior traces of the sigmoidal gene expression model. Genes with a large activation strength show a distinct gene expression pattern compared to those with a small activation strength. (**C**) A posterior density plot of the activation time for the five genes showing strong activation strength in (**B**).

Representative examples of genes with large and small activation strengths showed marked differences in the gene expression patterns corresponding to strong and weak switch-like behaviour as expected (Figure 7B). In addition, we examined the posterior density activation time *t*_0_ for the five genes showing strong switching behaviour (Figure 7C). Under a point estimate of pseudotime each gene would give a distinct activation time with which these genes can be ordered. However, when pseudotime uncertainty is taken into account, a distribution over possible activation times emerges. In this case, the five genes all have activation times between 0.3 and 0.5 precluding a precise ordering (if one exists) of activation. Visually, this seems sensible since there is considerable cell-to-cell variability in the expression of these genes and not all cells express the genes during the “on” phase. We are therefore unable to determine whether the “on” phase begins when the first cell with high expression is first observed in pseudotime or, if it starts before, and the first few cells simply have null expression (for biological or technical reasons).

We further explore this in Figure 8 which shows ten genes identified as having significant switch-like pseudotime dependence but with a range of mean activation times *t*_0_. The switchlike behaviour is stable to the different posterior pseudotime orderings that were sampled from the GPLVM. It is clear that the two genes RARRES3 and C1S are activating at an earlier time compared to the genes IL20RA and APOL4. However, we cannot be confident of the ordering within the pairs RARRES3/C1S and IL20RA/APOL4 in pseudotime since the distributions over the activation times are not well-separated and it is impossible to make any definitive statements as to whether one of these genes (in)activates before another. If the probability of a sequence of activation events is required, instead of examining each gene in isolation, we can count the number of posterior samples in which one gene precedes another instead and evidence may emerge of a possible ordering. These observations suggests a finite temporal resolution limit that can be obtained using pseudotemporal ordering.

**Figure 8:**
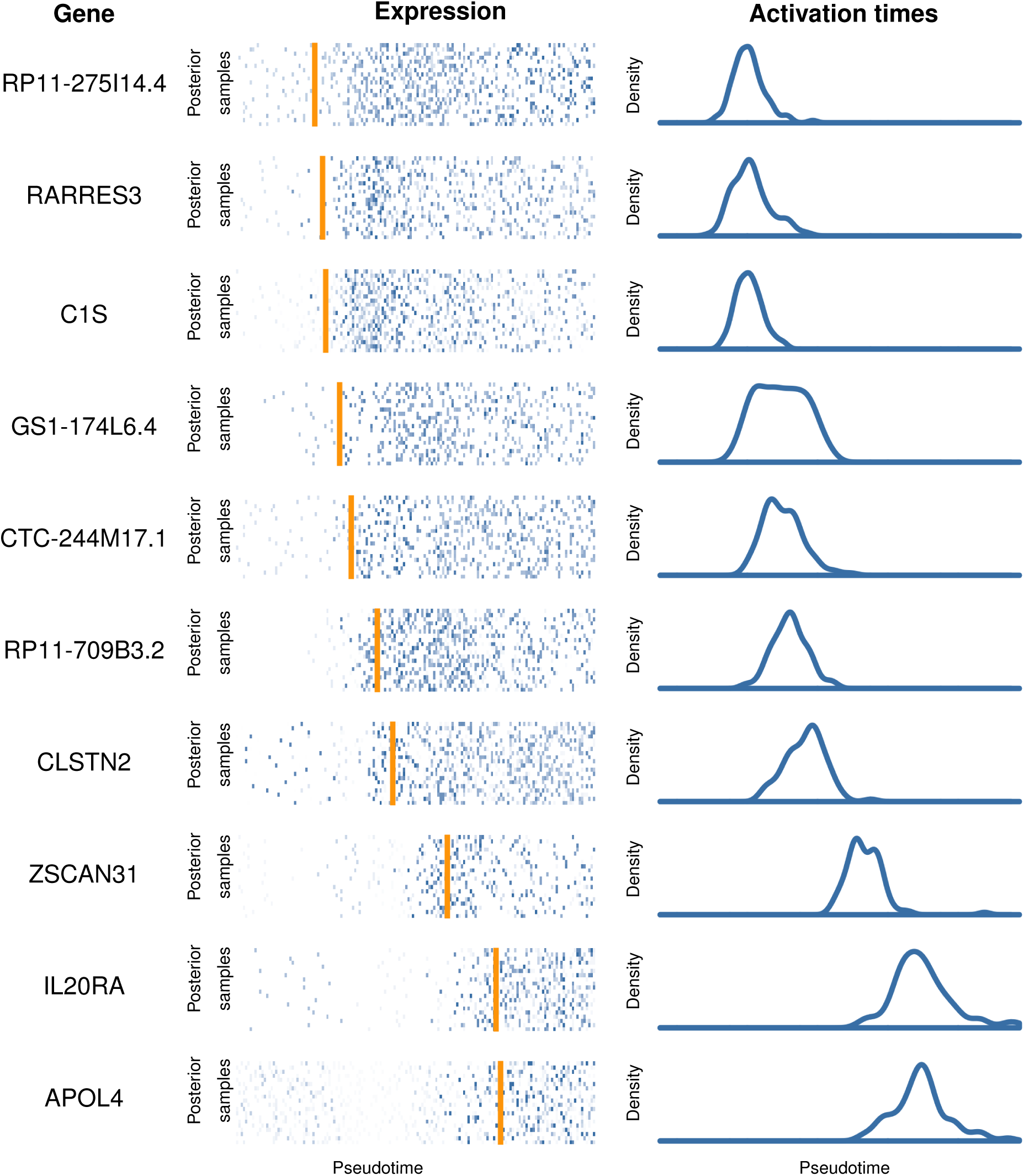
Identifying pseudotime dependent gene activation behaviour. Ten selected genes from Trapnell et al. (2014) found using our sigmoidal gene activation model exhibiting a range of activation times. For each gene, we show the expression levels of each cell (centre) where each row corresponds to an ordering according to a different posterior samples of pseudotime. The orange line corresponds to a point estimate of the activation time. The posterior density of the estimated activation time is also shown (right).

We note that we have deliberately avoid directly linking the sigmoidal gene activation and GPLVM pseudotime models to derive a single, joint model. In a joint model, the inference would attempt to order the cells in such a way as to maximise the fit of the sigmoidal and GPLVM to the expression data. However, as the sigmoidal model is only intended to identify genes with switch-like behaviour, it cannot explain other types of pseudotime dependence that may and do exist. This model misspecification would potentially drive inference in ways that cannot be foreseen.

### Learning trajectories from multiple reduced data representations

Finally, we address the impact of the subjective choice of dimensionality reduction that is normally applied to single cell gene expression data prior to pseudotime ordering and estimation. Typically, the choice of dimensionality reduction approach is based on whether the method gives rise to a putative pseudotime trajectory in the reduced dimensionality representation from visual inspection followed by confirmational analysis by examining known marker genes with established temporal association. This may lead to a number of possibilities since the same trajectory may exist in a number of reduced dimensionality representations.

One characteristic of the GPLVM is that the likelihood is conditionally independent across input dimensions. A consequence of this is that we can integrate heterogeneous data sources to learn pseudotimes as there is no requirement that each input dimension should come from the same representation or assay. We exploited this feature to examine the effect of the initial dimensionality reduction stage and see if we can learn pseudotime trajectories from multiple reduced dimension representations. Many dimensionality reduction algorithms have been applied to single-cell RNA-seq data, including PCA Shin et al. (2015), ICA Trapnell et al. (2014), t-SNE Marco et al. (2014) and diffusion maps Haghverdi et al. (2015).

We applied our probabilistic pseudotime inference algorithm to Laplacian Eigenmaps, PCA and t-SNE representations of the three datasets under consideration. We also applied the algorithm using all three representations jointly the results of which can be seen in Figure 9. While the pseudotime inference algorithm can fit trajectories to all three datasets individually, combining multiple representations can lead to a clearer, better defined trajectory. This allows us to track the same progression of cells through multiple reduced dimension representations at once, providing an equivalence to the trajectories represented by different dimensionality reduction algorithms. In fact there may be no correspondence between the trajectories in the different reduced dimension representations at all if the analysis is not integrated.

It should be noted that this approach is not necessarily an ideal integration model and more complex multi-view learning models Xu et al. (2013) should be investigated in future that will resolve the potential dependencies between the input representations. However, as a number of popular dimensionality reduction techniques (e.g. t-SNE) have no probabilistic interpretation and possess no underlying generative model, it is challenging to incorporate these within a coherent probabilistic framework (i.e. there is no likelihood function). The suggested technique, though implying simplistic independence assumptions, has practical value for incorporating such representations.

**Figure 9:**
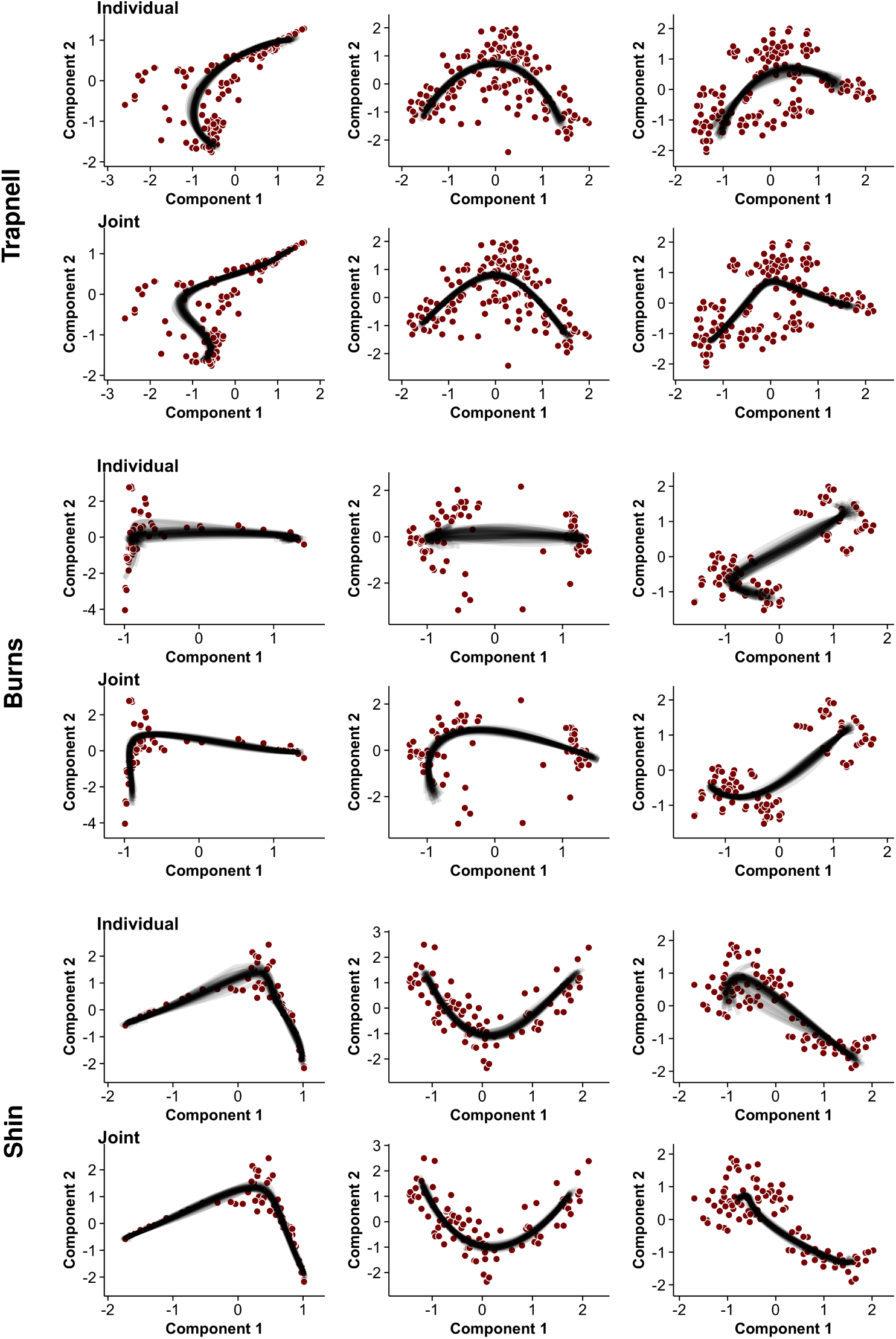
Learning pseudotime from three reduced dimension representations (Laplacian eigenmaps, PCA and t-SNE) of single-cell gene expression data from the three datasets studied (Trapnell, Burns and Shin). For each dataset the left column shows the Laplacian Eigenmaps representation, middle shows PCA and right shows t-SNE (Supplementary Methods). Pseudotime trajectories are fitted either on each representation individually (top row of each dataset) or jointly for all representations (bottom row). It can be seen that trajectory fits are more stable when the joint representations are used. Such analysis allows us to track cellular trajectories across multiple visualisations showing an equivalency of dimensionality reduction algorithms in the context of single-cell RNA-seq data.

We caution though that this integration approach is not intended to contain any arbitrary number of representations provided by the user. As each representation is ultimately drawn from the same underlying raw data, a representations should only be included if it provides some orthogonal (near-independent) information since the GPLVM assumes the representations are independent. In practice, this means selecting a small number of representations drawn from very different dimensionality reduction approaches. If the representations are not independent and are related this can give rise to an artificial reduction in posterior variance since we would be essentially doubling sample size by replicating the same data.

## Discussion

Pseudotime ordering from gene expression profiling of single cells provides the ability to extract temporal information about complex biological processes from which *true* time series experimentation may be technically challenging or impossible. In our investigations we have sought to characterise the utility of a probabilistic approach to the single cell pseudotime ordering problem over approaches that only return a single point estimate of pseudotime. Our work is significant since it has so far not been possible to assess the impact of this statistical uncertainty in downstream analyses and to ascertain the level of temporal resolution that can be obtained. In order to address this we adopted a Gaussian Process Latent Variable modelling framework to perform probabilistic pseudotime ordering within a Bayesian inference setting. The GPLVM allows us to probabilistically explore a range of different pseudotime trajectories within the reduced dimensional space. We showed that in a truly unbiased and unsupervised analysis the properties of the pseudotime trajectory will never purely be a product of the data alone and can heavily depend on prior assumptions about the smoothness, length scales of the trajectory and noise properties. Using samples drawn from the posterior distribution over pseudotime orderings under the GPLVM we were able to assess if genes that showed a significant pseudotime dependence under a point (MAP) pseudotime estimate would be robust to different possible pseudotime orderings. In two of the three datasets we examined we discovered that, when adjusted for pseudotime uncertainty, the false discovery rate may be significantly larger than the target 5%. Our investigations show that reliance on a single estimate of pseudotime ordering can lead to increased number of false discoveries but that it is possible to assess the impact of such assumptions within a probabilistic framework.

It is important to note that the GPLVM used in our investigations is not intended to be a single, all-encompassing solution for pseudotime modelling problems. For our purposes, it provided a simple and relevant device for tackling the single trajectory pseudotime problem in a probabilistic manner but clearly has limitations when the temporal process under investigation contains bifurcations or heteroscedastic noise processes (as discussed earlier). Improved and/or alternative probabilistic models are required to address more challenging modelling scenarios but the general procedures we describe are generic and should be applicable to any problem where statistical inference for a probabilistic model can give posterior simulation samples.

We also developed a novel sigmoidal gene expression temporal association model that enabled us to identify genes exhibiting a strong switch-like (in)activation behaviour. For these genes we were then able to estimate the activation times and use these to assess the time resolution that can be attained using pseudotime ordering of single cells. Our investigations show that pseudotime uncertainty prevents precise characterisation of the gene activation time but a probabilistic model can provide a distribution over the possibilities. In application, this uncertainty means that it is challenging to make precise statements about when regulatory factors will turn on or off and if they act in unison. This places an upper limit on the accuracy of dynamic gene regulation models and causal relationships between genes that could be built from the single cell expression data.

In conclusion, single cell genomics has provided a precision tool with which to interrogate complex temporal biological processes. However, as widely reported in recent studies, the properties of single cell gene expression data are complex and highly variable. We have shown that the many sources of variability can contribute to significant uncertainty in statistical inference for pseudotemporal ordering problems. We argue therefore that strong statistical foundations are vital and that probabilistic methods for provide a platform for quantifying uncertainty in pseudotemporal ordering which can be used to more robustly identify genes that are differentially expressed over time. Robust statistical procedures can also temper potentially unrealistic expectations about the level of temporal resolution that can be obtained from computationally-based pseudotime ordering. Ultimately, as the raw input data is not true time series data, pseudotime ordering is only ever an attempt to solve a *missing data* statistical inference problem that we should remind ourselves involves quantities (pseudotimes) that are *unknown, never can be known*.

## Methods

In addition to the descriptions below, further methodological descriptions and links to code to reproduce all our findings are given in Supplementary Methods.

### Statistical model for probabilistic pseudotime

The hierarchical model specification for the Gaussian Process Latent Variable model is described as follows:

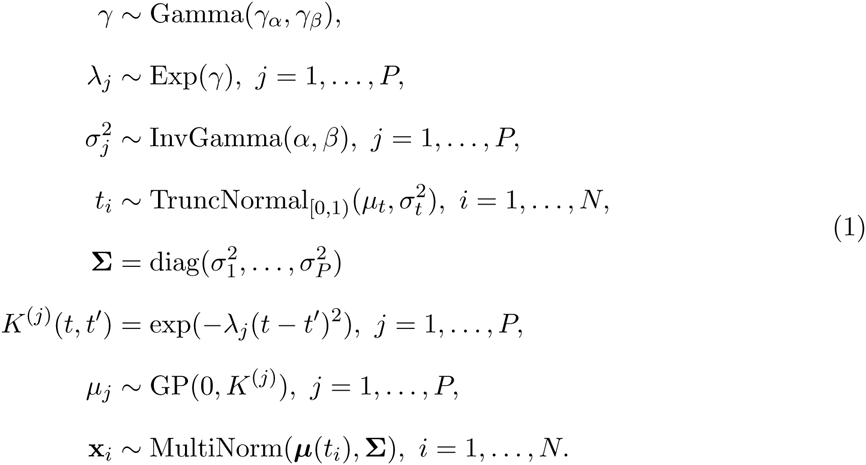

where x_i_ is the *P*-dimensional input of cell *i* (of *N*) found by performing dimensionality reduction on the entire gene set (for our experiments *P* = 2 following previous studies). The observed data is distributed according to a multivariate normal distribution with mean function **μ** and a diagonal noise covariance matrix **Σ**. The prior over the mean function **μ** in each dimension is given by a Gaussian Process with zero mean and covariance function *K* given by a standard double exponential kernel. The latent pseudotimes *t*_1_,…, *t*_*N*_ are drawn from a truncated Normal distribution on the range [0,1). Under this model |**λ**| can be thought of as the arc-length of the pseudotime trajectories, so applying larger levels of shrinkage to it will result in smoother trajectories passing through the point space. This shrinkage is ultimately controlled by the gamma hyperprior on *γ*, whose mean and variance are given by 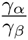 and 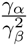 respectively. Therefore, adjusting these parameters allows curves to match prior smoothness expectations provided by plotting marker genes.

The hyperparameters *γ*_α_, *γ*_β_, *α*, *β*, *μ*_*t*_ and σ_*t*_^2^ are fixed and values for specific experiments for given in Supplementary Information. Inference was performed using the Stan probabilistic programming language Gelman et al. (2015) and our implementation is available as an **R** package at http://www.github.com/kieranrcampbell/pseudogp.

### Integrating multiple representations

One feature of the GPLVM is that the likelihood is conditionally independent (given the pseudotimes) across input dimensions. If we have a set of *Q* reduced dimension representations of single-cell data {**X**_*i*_, *i* = 1,…, *Q*} (be they multiple representations of the same assay, e.g. RNA-seq, or multiple representations of multiple assays) then the likelihood factorises across each representation. For example, if we have Laplacian Eigenmaps, PCA and t-SNE representations of the same data then the likelihood becomes

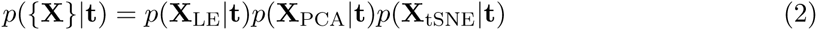

where **t** is the pseudotime vector to be learned and inference proceeds straightforwardly using this product likelihood.

### Sigmoidal model for switch-like gene (in)activation behaviours across pseudo-time

We detail the mathematical specification of the sigmoidal switch model below. Let *y*_*ij*_ debotes the log_2_ gene expression of gene *i* in cell *j* at pseudotime *t*_*j*_ then

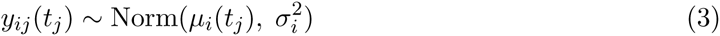

where

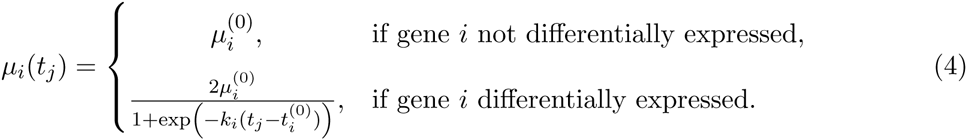

Under this model the parameter *k*_*i*_ can be thought of as an activation ‘strength’ relating to how quickly a gene switches on or off, while 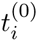 relates to the pseudotime at which the gene switches on or off.

The case of a gene not being differentially expressed is a nested model of the differential expression case found by setting *k* = 0. Consequently we can use a likelihood ratio test with no differential expression as the null hypothesis and differential expression as the alternative and twice the difference in their log-likelihoods will form a *χ*^2^ test statistic with 2 degrees of freedom. The maximum likelihood estimates of the parameters under the differential expression model have no analytical solution so L-BFGS-B optimisation was used (implemented in the **R** package switchde, http://github.com/kieranrcampbell/switchde).

## Author’s contributions

K.C. and C.Y. conceived the study. K.C. developed software and performed computer simulations. K.C. and C.Y. wrote the manuscript.

## Acknowledgements

K.C. is supported by a UK Medical Research Council funded doctoral studentship. C.Y. is supported by a UK Medical Research Council New Investigator Research Grant (Ref. No. MR/L001411/1), the Wellcome Trust Core Award Grant Number 090532/Z/09/Z, the John Fell Oxford University Press (OUP) Research Fund and the Li Ka Shing Foundation via a Oxford-Stanford Big Data in Human Health Seed Grant.

## References

1. Amir, E.-a. D., K. L. Davis, M. D. Tadmor, E. F. Simonds, J. H. Levine, S. C. Bendall, D. K. Shenfeld, S. Krishnaswamy, G. P. Nolan, and D. Pe’er (2013, June). viSNE enables visualization of high dimensional single-cell data and reveals phenotypic heterogeneity of leukemia. Nature biotechnology 31(6), 545–52.

2. Belkin, M. and P. Niyogi (2003). Laplacian Eigenmaps for Dimensionality Reduction and Data. 1396, 1373–1396.

3. Bendall, S. C., K. L. Davis, E.-A. D. Amir, M. D. Tadmor, E. F. Simonds, T. J. Chen, D. K. Shenfeld, G. P. Nolan, and D. Pe’er (2014, April). Single-cell trajectory detection uncovers progression and regulatory coordination in human B cell development. Cell 31(6), 545–52.

4. Burns, J. C., M. C. Kelly, M. Hoa, R. J. Morell, and M. W. Kelley (2015). Single-cell RNA-Seq resolves cellular complexity in sensory organs from the neonatal inner ear. Nature Communications 6, 8557.

5. Campbell, K. and C. Yau (2015). Bayesian gaussian process latent variable models for pseudo-time inference in single-cell rna-seq data. bioRxiv, 026872.

6. Gelman, A., D. Lee, and J. Guo (2015). Stan a probabilistic programming language for bayesian inference and optimization. Journal of Educational and Behavioral Statistics, 1076998615606113.

7. Gupta, A. and Z. Bar-Joseph (2008). Extracting dynamics from static cancer expression data. IEEE/ACM transactions on computational biology and bioinformatics / IEEE, ACM 31(6), 545–52.

8. Haghverdi, L., F. Buettner, and F. J. Theis (2015). Diffusion maps for high-dimensional single-cell analysis of differentiation data. Bioinformatics (May), 1–10.

9. Hastie, T. and W. Stuetzle (2012, March). Principal Curves.

10. Hinton, G. E. and S. T. Roweis (2002). Stochastic neighbor embedding. In Advances in neural information processing systems, pp. 833–840.

11. Kalisky, T. and S. R. Quake (2011). Single-cell genomics. Nature methods 31(6), 545–52.

12. Le, Q. V., A. J. Smola, and S. Canu (2005). Heteroscedastic gaussian process regression. In Proceedings of the 22nd international conference on Machine learning, pp. 489–496. ACM.

13. Macaulay, I. C. and T. Voet (2014, January). Single cell genomics: advances and future perspectives. PLoS genetics 10(1), e1004126.

14. Magwene, P. M., P. Lizardi, and J. Kim (2003). Reconstructing the temporal ordering of biological samples using microarray data. Bioinformatics 31(6), 545–52.

15. Marco, E., R. L. Karp, G. Guo, P. Robson, A. H. Hart, L. Trippa, and G.-C. Yuan (2014, December). Bifurcation analysis of single-cell gene expression data reveals epigenetic landscape. Proceedings of the National Academy of Sciences of the United States of America 111(52), E5643–50.

16. Moignard, V., S. Woodhouse, L. Haghverdi, A. J. Lilly, Y. Tanaka, A. C. Wilkinson, F. Buet-tner, I. C. Macaulay, W. Jawaid, E. Diamanti, S.-I. Nishikawa, N. Piterman, V. Kouskoff, F. J. Theis, J. Fisher, and B. Gttgens (2015, February). Decoding the regulatory network of early blood development from single-cell gene expression measurements. Nature Biotechnology 33(3).

17. Qiu, P., A. J. Gentles, and S. K. Plevritis (2011, April). Discovering biological progression underlying microarray samples. PLoS computational biology 7(4), e1001123.

18. Qiu, P., E. F. Simonds, S. C. Bendall, K. D. Gibbs Jr, R. V. Bruggner, M. D. Linderman, K. Sachs, G. P. Nolan, and S. K. Plevritis (2011). Extracting a cellular hierarchy from high-dimensional cytometry data with spade. Nature biotechnology 31(6), 545–52.

19. Reid, J. E. and L. Wernisch (2015). Pseudotime estimation: deconfounding single cell time series. bioRxiv, 019588.

20. Shapiro, E., T. Biezuner, and S. Linnarsson (2013, September). Single-cell sequencing-based technologies will revolutionize whole-organism science. Nature reviews. Genetics 14(9), 618–30.

21. Shin, J., D. A. Berg, Y. Zhu, J. Y. Shin, J. Song, M. A. Bonaguidi, G. Enikolopov, D. W. Nauen, K. M. Christian, G.-l. Ming, and H. Song (2015, August). Single-Cell RNA-Seq with Waterfall Reveals Molecular Cascades underlying Adult Neurogenesis. Cell Stem Cell 31(6), 545–52.

22. Stegle, O., S. a. Teichmann, and J. C. Marioni (2015, January). Computational and analytical challenges in single-cell transcriptomics. Nature Reviews Genetics 31(6), 545–52.

23. Titsias, M. and N. Lawrence (2010). Bayesian Gaussian Process Latent Variable Model. Artificial Intelligence 9, 844–851.

24. Trapnell, C. (2015, Oct). Defining cell types and states with single-cell genomics. Genome Res 31(6), 545–52.

25. Trapnell, C., D. Cacchiarelli, J. Grimsby, P. Pokharel, S. Li, M. Morse, N. J. Lennon, K. J. Livak, T. S. Mikkelsen, and J. L. Rinn (2014, April). The dynamics and regulators of cell fate decisions are revealed by pseudotemporal ordering of single cells. Nature biotechnology 31(6), 545–52.

26. Treutlein, B., D. G. Brownfield, A. R. Wu, N. F. Neff, G. L. Mantalas, F. H. Espinoza, T. J. Desai, M. A. Krasnow, and S. R. Quake (2014). Reconstructing lineage hierarchies of the distal lung epithelium using single-cell rna-seq. Nature 31(6), 545–52.

27. Tsang, J. C., Y. Yu, S. Burke, F. Buettner, C. Wang, A. A. Kolodziejczyk, S. A. Teichmann, L. Lu, and P. Liu (2015). Single-cell transcriptomic reconstruction reveals cell cycle and multi-lineage differentiation defects in bcl11a-deficient hematopoietic stem cells. Genome biology 31(6), 545–52.

28. Van der Maaten, L. and G. Hinton (2008). Visualizing data using t-sne. Journal of Machine Learning Research 9(2579-2605), 85.

29. Wills, Q. F. and A. J. Mead (2015). Application of single cell genomics in cancer: Promise and challenges. Human molecular genetics, ddv235.

30. Xu, C., D. Tao, and C. Xu (2013). A survey on multi-view learning. arXiv preprint arXiv:1304.5634.

